# Sex-stratified insights into the genetics of brain volumes in late adulthood

**DOI:** 10.64898/2026.01.09.698629

**Authors:** Édith Breton, Elleke Tissink, Gloria Matte Bon, Dennis van der Meer, Ole A. Andreassen, Tobias Kaufmann

**Affiliations:** Centre for Precision Psychiatry, Division of Mental Health and Addiction, Institute of Clinical Medicine, University of Oslo, Oslo, Norway; Centre intersectoriel en santé durable, département des sciences fondamentales, Université du Québec à Chicoutimi, Saguenay, Canada; Centre intégré universitaire de santé et de services sociaux (CIUSSS) du Saguenay-Lac-Saint-Jean, Saguenay, QC, Canada; Department of Psychiatry and Psychotherapy, Tübingen Center for Mental Health, University of Tübingen, Tübingen, Germany; Department of Women’s and Children’s Health, Science for Life Laboratory, Uppsala University, Uppsala, Sweden; German Center for Mental Health (DZPG), partner site Tübingen, Tübingen, Germany

**Keywords:** Sex Differences, Brain Anatomy, Imaging Genetics

## Abstract

**Background:** The prevalence, timing and disease course of mental and neurological disorders vary according to sex, yet the neurogenetic mechanisms underlying sex differences in these disorders remain poorly understood. *<u>Methods.</u>* Here, we explored the genetic architecture of the anatomical volumes of 257 regions of interest across the brain using multivariate genome-wide analyses in 15,740 male and 15,740 female participants from the UK Biobank, matched on age and scan site.

**Results:** Our findings revealed that the genetics of brain volume is highly similar between females and males in late adulthood. Yet, we found evidence of possible autosomal sex heterogeneity, particularly in the number of brain-volume associated genes, which was higher in females than in males. Variability in the number of identified genes were marked in limbic regions such as the insula, the cingulate cortex, the hippocampus and the amygdala.

**Conclusion.:** Overall, our findings contribute to a better understanding of the genetic determinants of brain volumes in males and females. Because neurogenetics may also influence the risk for sex-prevalent brain disorders, the current findings have the potential to facilitate precision medicine approaches in improving prevention strategies and targeted treatments.

**Plain English summary:** Mental and neurobiological disorders often differ between females and males. Some disorders are more common in one sex than the other, and symptoms can present or evolve differently. Variation in brain biology, partly shaped by genetic factors, may contribute to these observed patterns. In this study, we investigated genetic factors linked to brain structure in females and males separately. We focused on the volumes of 257 brain regions and analyzed genetic data from more than 30,000 adults from the UK Biobank. Overall, we found that the patterns of association between genetics and brain volumes are largely similar between females and males in late adulthood. At the same time, we observed evidence of sex-dependent patterns, particularly in the number of genes associated with brain volume, which tended to be higher in females than in males. These differences were marked in brain regions involved in the limbic system. Together, our findings improve our understanding of how genetics contribute to brain structure in females and males.

**Highlights:** Genetic influences on brain volumes are largely shared between females and males in late adulthood.

Gene-level variability was observed, with a higher number of brain volumes associated genes identified in females than in male, in both multivariate and univariate analyses.

This gene-level variability was most pronounced in limbic regions, such as the insular and cingulate cortices.

## Background

Sex impacts the prevalence, clinical presentation, course and treatment response of several mental and neurological disorders (1,2). Major depressive disorder and Alzheimer’s disease are twice as common in females as in males, and anorexia nervosa is 10 times more common in females than in males (3–5). In contrast, autism spectrum disorder, alcohol use disorder and attention deficit hyperactivity disorder are two to three times more common in males (6–8). The main windows of vulnerability for mental disorders also differ by sex, with females more likely to be affected by conditions that begin in adolescence or later, and males more likely to develop neurodevelopmental disorders that express in childhood. Yet, the biological mechanisms underlying these differences have not been elucidated (9,10). Such mechanistic understanding could facilitate efforts towards precision medicine in the clinical neurosciences and contribute to reducing sex inequalities in health care and research (11,12).

Neuroimaging studies have identified structural and functional brain differences between males and females across the lifespan, including greater variability in brain structure in males (13). There is evidence to suggest that these brain differences may be associated with mental disorders (14), but more studies are needed to establish this association. Sex chromosomes and hormones are obvious contributors to the sex differences in the brain (15), but they are not the only ones. For example, genetic studies have identified sex-dependent associations between autosomal gene variants and mood or psychotic disorders (16). There is also evidence that autosomal gene variants interact with sex hormones to differentially affect the risk of conditions such as dementia (17).

Yet, there is still little understanding of the interaction between sex, genetics and the brain, and few large-scale studies have considered study designs that analyze females and males separately (12). Existing evidence suggests that common genetic variants often impact gene expression differently in the brains of males and females (18), which may be relevant for disorder risk (19). Further studies have suggested that genes involved in neurotransmission, immune function and mental disorders are differently targeted by transcriptional regulators in both sexes (20). Throughout the life course, epigenetic and other regulatory factors, inflammatory processes, neurodevelopmental processes and ageing all plausibly contribute to shaping the genetics-brain interaction differently in males and females, potentially influencing their vulnerability to common brain disorders.

Here, we investigated the genetic architecture of brain volume in 15,740 females and 15,740 males from the UK Biobank. Both groups were matched for age and scan site. Our overarching aim was to gain further insight into how genetics are linked to brain structure in females and males. We opted for sex-stratified analyses to characterize whether the overall patterns of autosomal genetic associations observed in one sex also show up in the other. To this end, we extracted volumes of 180 cortical and 77 subcortical regions across the entire brain for genome-wide association studies (GWAS). We leveraged MOSTest (21), a multivariate GWAS tool designed to enhance power and facilitate discovery in imaging genetics research, yielding one multivariate summary statistic across the whole brain for females and males separately. In addition, we assessed each brain region in females and males separately using a univariate approach.

## Methods

### Sample

We selected participants from the UK Biobank with genetic and neuroimaging data and of European descent, defined by self-identification as “White British” and similar genetic ancestry based on genetic principal components analysis (accession code no. 27412). Study procedures were approved by local institutional review boards and each sample was collected with participants’ written informed consent. Standardized brain MRI and genomic protocols are described elsewhere (22–24).

### Magnetic Resonance Imaging

The present study used T1-weighted magnetic imaging volumes extracted using FreeSurfer (https://surfer.nmr.mgh.harvard.edu/). The whole brain volume measure correspond to the mean of the right and left hemisphere volumes, and included a combination of 180 cortical ROIs based on the Glasser Atlas (25), as well as subcortical structures (10 ROIs (26)) and further segmentation of the amygdala (9 nuclei + whole amygdala (27)), the hippocampus (19 subregions + whole hippocampus (28)), the thalamus (25 subregions + whole thalamus (29)), the brainstem (4 subregions + whole brainstem (30)) and the hypothalamus (5 subregions + whole hypothalamus (31)), for a total of 257 ROIs.

Individual structures with volumes exceeding three standard deviations below the mean of the Euler number were excluded from the analyses. This left us with 34,706 participants of European descent with genetics and neuroimaging data. From these 34,704 participants, we selected male-female pairs, ensuring matching for scan site and age (within a 6-months difference) while minimizing discrepancies in estimated total intracranial volume by selecting the pairs with the smallest differences, based on a matching procedure deployed by others (32). We made sure that there was no significant difference (*p*>0.05) between males and females for age and total intracranial volume, for each scan site. Participants who did not meet these match criteria were excluded, leading to the inclusion of 15,740 males and 15,740 females with a mean age of 64,4 years.

### Genome-wide analyses

We deployed the Multivariate Omnibus Statistical Test (MOSTest) (21) to perform two multivariate genome-wide association analyses of the whole brain volume, in females and males separately, focusing on autosomes. Individuals with genotype missing rate >10%, variants with genotype missing rate > 5%, variants failing Hardy-Weinberg equilibrium at *p*=1x10^-9^, variants with minor allele frequency below 0.5%, and imputation info score below 0.5 were filtered out. We controlled our analyses for covariates by pre-residualizing the ROI volumes for age, age^2^, scanning site, the first 20 genetic principal components, Euler number, and estimated total intracranial volume (in R version 4.2).

In parallel, we extracted univariate GWASs from the univariate stream of MOSTest (21) for each of the 257 ROIs, to gain a better understanding of the regional heterogeneity that may occur between males and females in specific brain regions.

### FUMA

The Functional Mapping and Annotation of Genome-Wide Association Studies (FUMA) platform (33) was used to perform positional gene mapping for all candidate SNPs, using the default settings of the SNP2GENE functionality. For the list of overlapping genes between females and males, FUMA GENE2FUNC was used to perform enrichment analyses against 15 496 gene sets (curated gene sets: 5 500, Gene Ontology terms: 9 996) from MsigDB version v7.0. We corrected for multiple testing using FDR and set the minimum overlapping genes to 2.

### Heritability & Genetic correlations

The sex-stratified univariate GWASs summary statistics were used to compute heritability and genetic correlations estimates for each ROI between females and males, using linkage disequilibrium score regression (LDSC) (34,35) with default settings and LD Scores based on the European 1000 Genomes reference panel. We filtered out phenotypes with low heritability z-scores (*h*^2^_SNP_ / SE < 1.96) from further analyses.

## Results

### The genetics of brain volumes is similar in females and males, with some variability at the gene level

Using a multivariate GWAS framework, we identified 231 genome-wide significant loci (*p* < 5e-8) associated with brain volumes in females and 241 loci associated with brain volumes in males, including 127 physically overlapping loci between both sexes (based on chromosomes, loci start positions and end positions). Genome-wide patterns of association appeared highly similar between sexes (**Figure 1A**). Among the identified loci, 199 (female) and 190 (male) loci could positionally be mapped to genes, leading to the identification of 1000 genes associated with brain volumes in females and 849 genes associated with brain volumes in males (since some loci map to many genes; **Figure 1B**). Approximately half of the identified genes (N=552) overlapped between males and females, while 448 genes were only found in females and 297 genes were only found in males (**Figure 1C**, **Supplementary Table 1)**.

**Figure 1:**
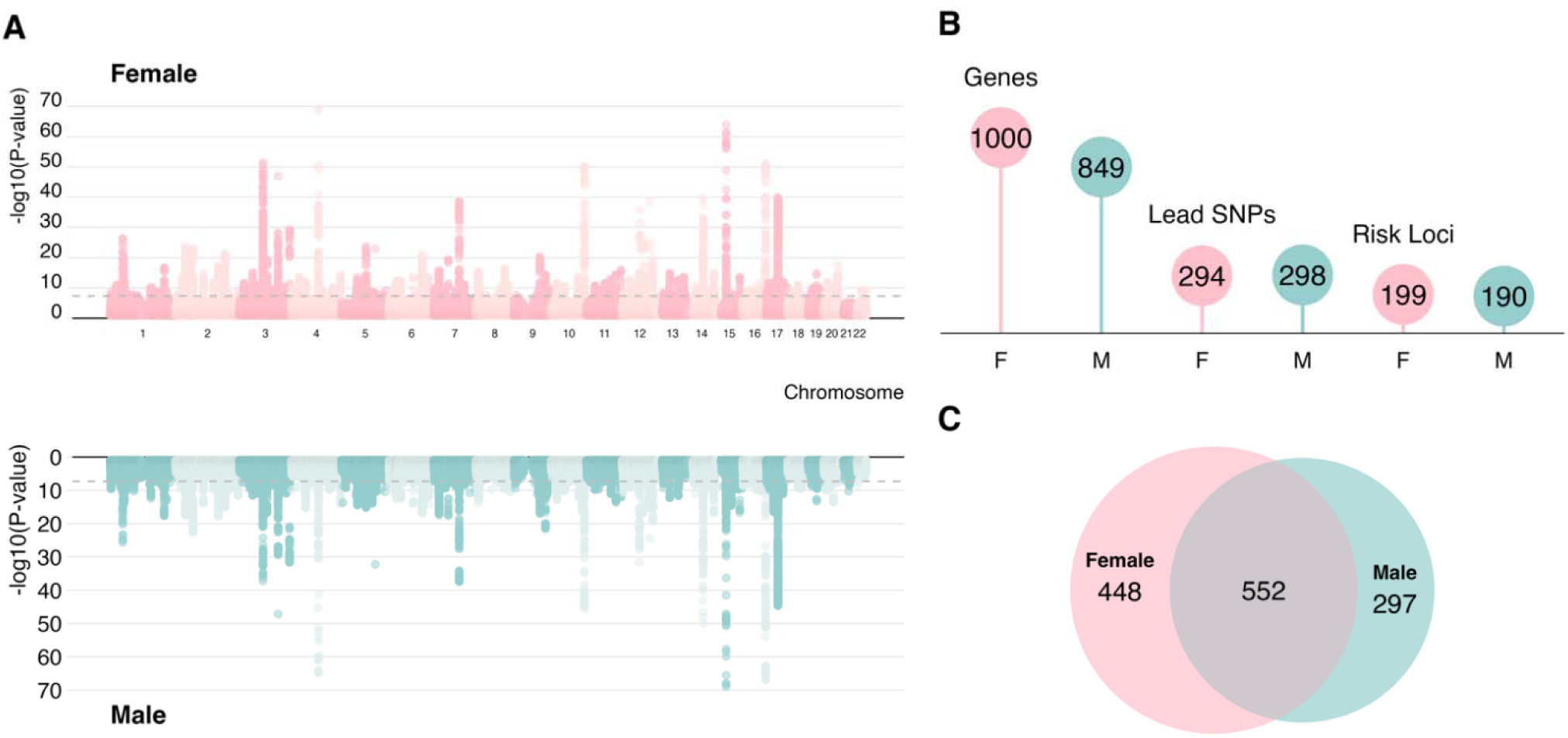
Sex-stratified multivariate analyses reveal shared and distinct genetic contributions to brain volume (A) Miami plot displaying the multivariate genome-wide association statistics in females (top/pink) and males (bottom/blue). These statistics reflect the association across 257 brain regions of interest. (B) Comparison of the number of genes, lead single nucleotide polymorphisms (SNPs) and risk loci identified in females (pink) versus males (blue) based on the multivariate GWAS statistics. The number of risk loci correspond to the number of loci with at least one gene mapped. (C) Venn diagram showing the corresponding number of genes overlapping or not in females (pink) and males (blue).

The genes shared between females and males were significantly (*p_FDR_* < 0.05) enriched in 123 gene-sets (see Methods, **Supplementary Figure 1**). 25 of these were related to neurodevelopmental pathways (e.g., neuron development, neurogenesis, regulation of neuron projection), 15 were related to developmental/growth processes, and 15 were related to immune pathways. Overlapping genes between males and females were significantly (*p_FDR_*< 0.05) enriched for differentially expressed gene-sets in 21 tissue types. These included 11 key brain regions such as the putamen, nucleus accumbens, caudate nucleus, amygdala, anterior cingulate cortex, hypothalamus and hippocampus, frontal cortex, and substantia nigra (**Supplementary Figure 2**).

The 448 genes only significantly associated with brain volumes in females were enriched in 23 tissue-specific gene-sets with differential expression, of which 13 were brain tissues (**Supplementary Figure 3**). Most of these (N=19) overlapped with the enrichment pattern of genes shared between females and males described above, except for cerebellum, cerebellar hemisphere, tibial artery and EBV-transformed lymphocytes.

On the other hand, genes only found as significant in males (N=297) were enriched for 34 differentially expressed tissue-specific gene-sets. These included 12 brain tissues that were also identified in genes only found for females or genes shared between males and females (**Supplementary Figure 4**).

### Divergences in number of genes identified in males versus females are widespread across the brain

Next, we sought to characterize patterns of sex-stratified associations across distinct brain regions. To do this, we turned to a univariate framework and performed standard GWASs for each of the 257 regions of interest (ROI) separately. We further assigned cortical ROIs to their respective lobes (i.e., frontal, parietal, temporal, occipital, insula, cingulate). We performed sex-stratified LDSC (34,35) to estimate SNP-based heritability across the 257 brain regions in females and males, yielding highly similar mean heritability estimates of h^2^=0.19 ± 0.07 in females and h^2^=0.18 ± 0.06 in males (**Supplementary Figure 5**). As expected, the genetic correlations across the genome were very high between the sexes in all cortical and subcortical brain regions (based on LDSC; all r >0.72; **Figure 2A-B; 4**). However, at the gene level, these regions showed potential sex-related heterogeneity. (**Figure 2C**).

**Figure 2:**
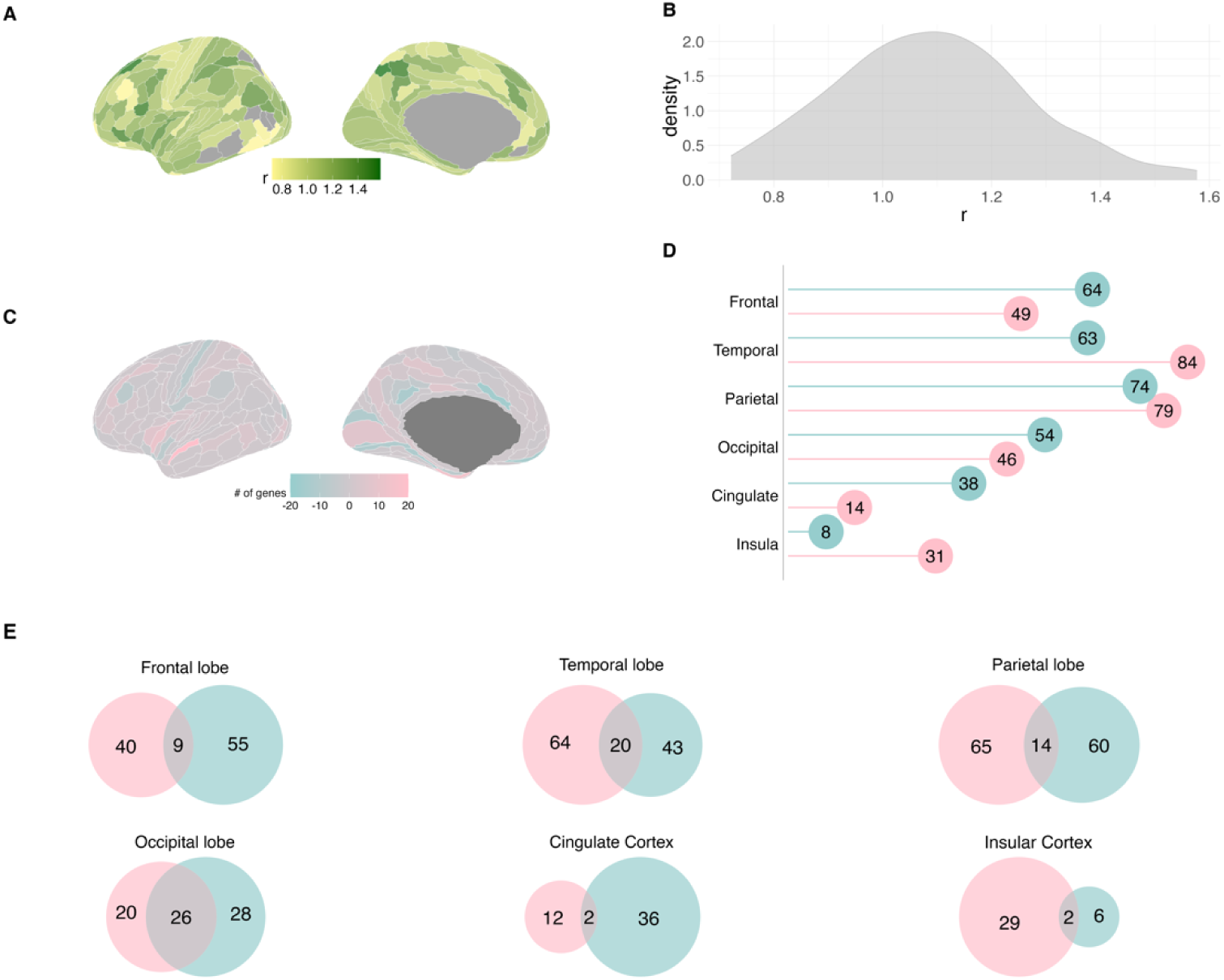
The genetic architecture of brain volume shows global similarity between sexes, yet exhibits divergent gene-level patterns (A) Genetic correlations for each cortical ROI between males and females. Note that LDSC correlation estimates are not bound to 1 and can therefore exceed 1 in cases of high GWAS similarity and depending on the estimation error. (B) Density plot showing the distribution of genetic correlations between males and females for all cortical ROIs. (C) Differences between females and males (i.e., females – males) in the number of genes identified through univariate GWASs, for each cortical regions of interest. Pink shades indicate a higher number of genes in females, whereas blue shades indicate a higher number of genes in males. (D) Number of genes identified in females (pink) versus males (blue) in the 180 univariate GWASs for cortical ROIs, presented by brain lobe. (E) Venn diagrams showing the overlap of genes identified in females (pink) and males (blue) across brain lobes, based on univariate GWASs. Full gene lists can be found in S2-S7.

Next, we examined sex-stratified patterns at the lobe level (**Figure 2D)**. Notably, the insular and cingulate cortices showed indications of greater sex heterogeneity. Particularly, the cingulate cortex was significantly associated to twice as many genes in males versus females, with 38 genes identified compared to 14 in females, with only 2 genes overlapping (**Figure 2E)**. In contrast, the insula was significantly associated with almost four times more genes in females (31 genes) than in males (8 genes), with only two genes overlapping between the sexes (**Figure 2E**).

In addition, we examined the overlap of genes identified by univariate GWASs in the different brain regions, both within and between sexes. Our analysis revealed limited gene overlap across brain regions and within and between sexes, highlighting regional and potential sex-dependent variability at the gene level (**Figure 3**).

**Figure 3:**
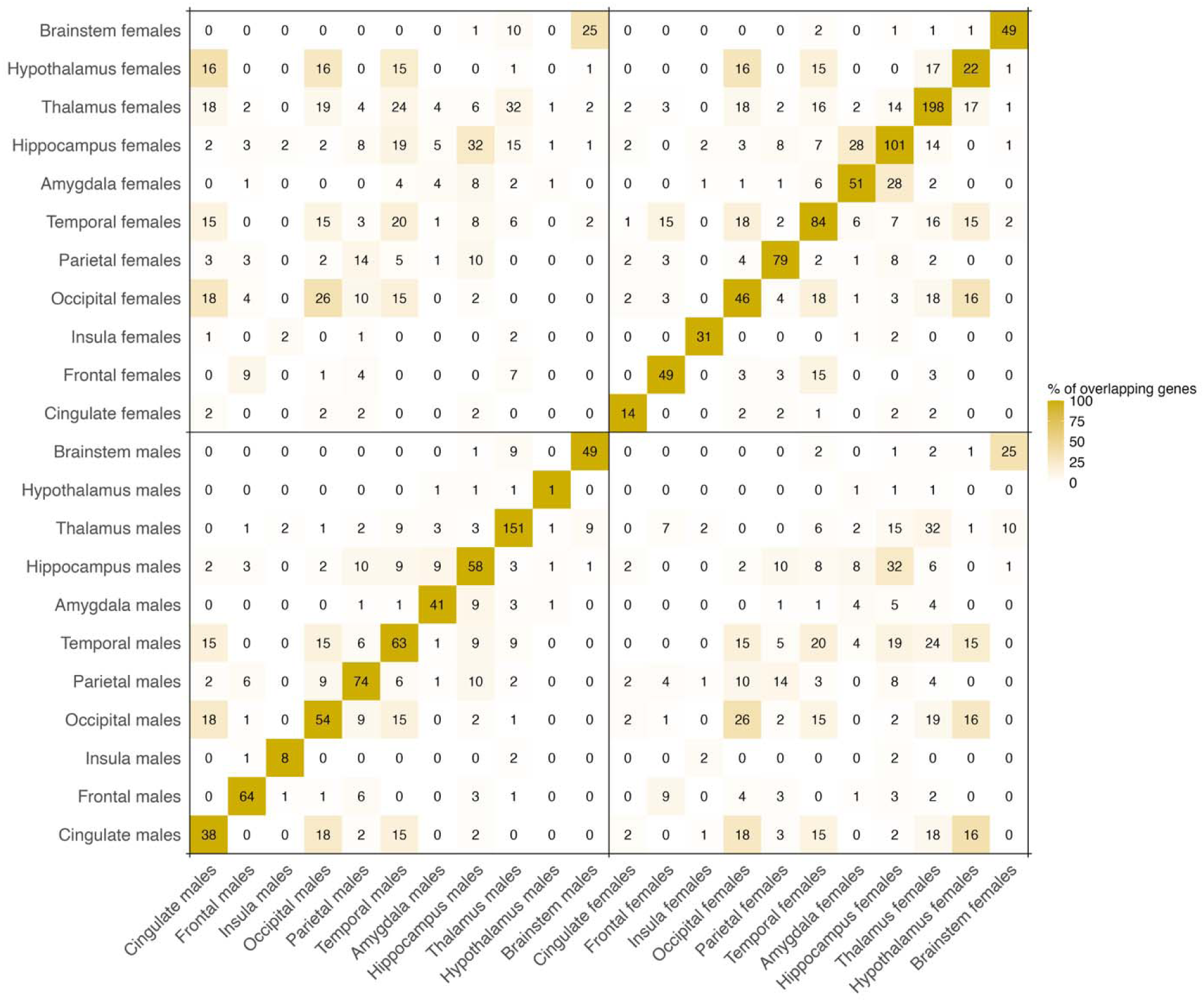
Genes identified by univariate analyses vary across brain regions and sexes Heatmap showing the percentage of overlapping genes out of the total number of genes identified for two given regions, for all cortical and subcortical brain regions, and within or between sexes. Color scale represents the percentage of overlapping genes between two regions whereas the absolute number of overlapping gene is displayed within each cell. The figure is based on univariate GWAS results. Full gene lists can be found in Supplementary Tables 2-12.

### Subcortical structures appear to be associated with more genes in females than in males

In line with the cortical results, univariate GWAS for subcortical regions were also strongly correlated between males and females (**Figure 4A**). All subcortical structures (except the brainstem) showed a higher number of genes in females than in males (**Figure 4B**).

**Figure 4:**
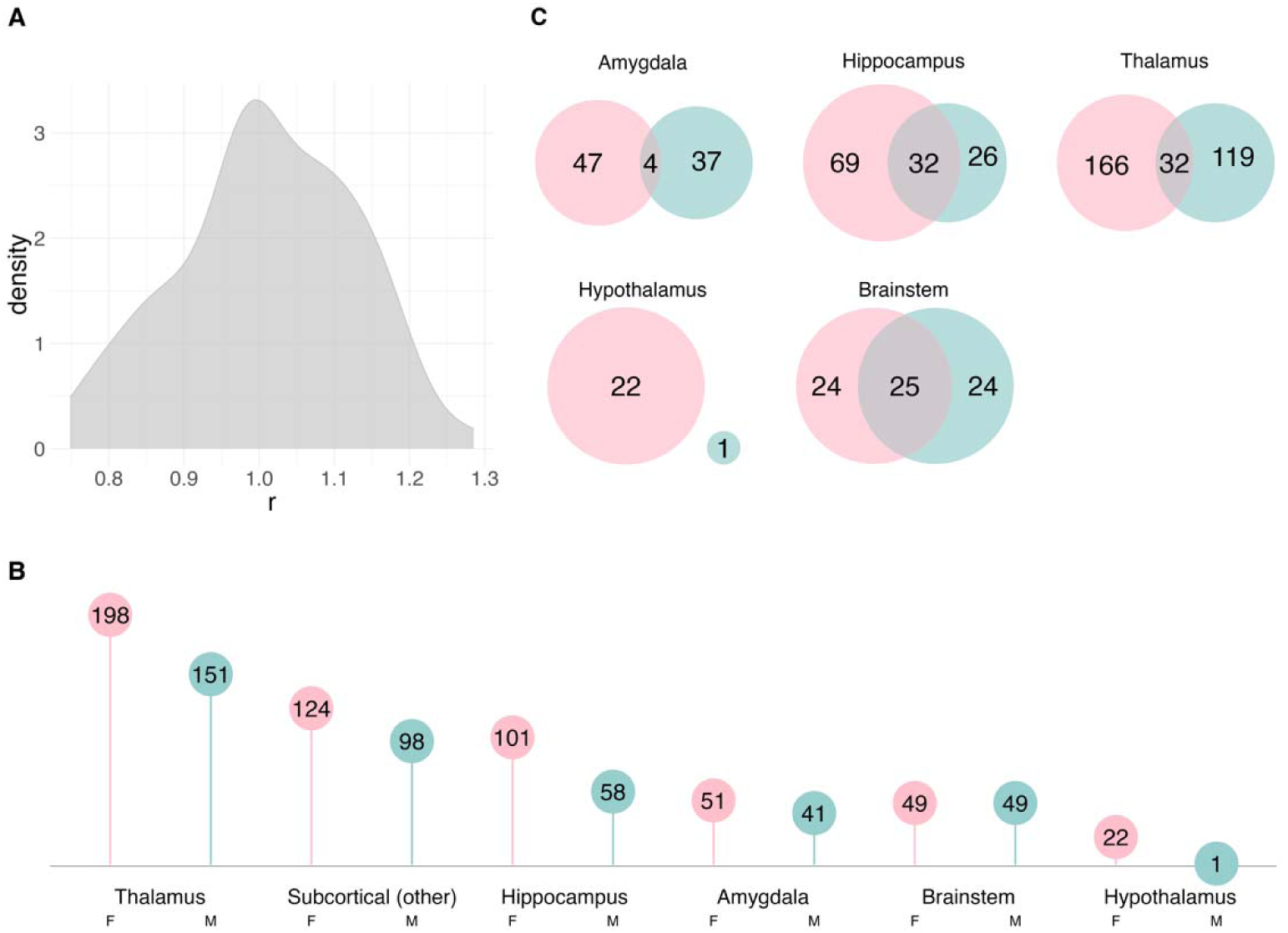
Limited overlap between females and males for genes associated with subcortical structures (A) Density plot showing the distribution of genetic correlations between males and females for all subcortical ROIs. (B) Comparison of the number of genes identified in females (pink) versus males (blue) by the 77 univariate GWASs for each subcortical ROI, presented by subcortical structure. (C) Venn Diagrams showing the number of genes identified by univariate GWASs in females (pink) and males (blue); for all ROIs located in subcortical structures. Full gene lists can be found in S8-S12.

Different genes emerged in females and males within limbic structures such as the amygdala and the hippocampus (**Figure 4C**; **Supplementary Tables 8-9)**. In the hippocampus, we identified 69 genes only genome-wide significant (*p* < 5e-8) in females and 26 genes only significant in males. In the amygdala, we found 47 genes in females, 37 genes in males and 4 overlapping genes. In the thalamus, only 32 genes were significant in both sex-stratified GWAS, whereas we identified 166 reached genome-wide significance in the female-stratified GWAS only, and 119 in the male-stratified GWAS only (**Figure 4C**, **Supplementary Table 10)**.

Lastly, there was no overlap between female and male genes for the hypothalamus (**Figure 4C**, **Supplementary Table 11)**.

## Discussion

The present study used a combination of multivariate and univariate genome-wide approaches to quantify and overlap the genes significantly associated with human brain anatomy in females and males separately. Genetic correlations between the sexes were high across all brain regions, indicating largely shared genetic architecture. Yet, we observed sets of genes that reached genome-wide significance in only one sex, possibly highlighting candidate regions for sex-dependent genetic effects. Moreover, our results suggest that a larger set of genes shows significant associations with brain volume anatomy in females than in males, at least in late adulthood. This excess of genome-wide significant genes in females aligns with recent reports demonstrating a higher number of genomic loci with significant local heritability in females across a wide range of quantitative traits (36).

We found that, particularly in limbic structures, different genes were significant for females versus males, with many structures being significantly associated with a higher number of genes in females compared to males. Responsible for emotions, learning and memory (37), the limbic system has been linked to variation between males and females, both in terms of brain structure and association with neurological and psychiatric disorders. A previous study by our group found that limbic structures show greater volume differences between sexes than non-limbic structures, and that they are more heritable (38). In the present study, the cingulate cortex was the only region where more genes were identified in males. Sex differences in the cingulate cortex have been reported in healthy individuals (39), but also in people with autism spectrum disorders (40) and depression (41). The cingulate cortex is also associated with impulsivity and aggressive behaviors in childhood, and these associations have been shown to differ between males and females (42,43). In line with these evidence, most of the cingulate genes only identified as significant in males have previously been associated with male-prevalent mental disorders, including autism spectrum disorders (e.g., *RPP25* (44)) and cocaine dependence (e.g., *NSF* (45)).

The present study identified genes associated with brain volumes that could potentially be linked to variations in the clinical presentation of common brain disorders, although further research is needed to confirm this hypothesis. As an example, in the hippocampus and thalamus of females—but not males—we respectively found *CARF*, *NBEAL1*, and *LRP11*, as well as *CRHR1*, *NSUN3*, and *NCKIPSD*. These genes have previously been implicated in eating disorders or anxiety – conditions that disproportionately affect females (46,47). In contrast, the genes that we identified as significant only in males were frequently involved in male-prevalent conditions that typically onset in childhood, such as autism spectrum disorders, intellectual disability and epilepsy (e.g., *AUTS2* (48), *RPP25* (44), *SLC1A2* (49)). Overall, our findings are a first step in exploring how neurogenetic variation could relate to behavioural and clinical sex differences in brain disorders. Sex-related variability has long been overlooked in research; yet, a deeper understanding of the underlying biological mechanisms is crucial to translate neuroscientific advances into clinical applications that can meaningfully reduce mental health inequalities (1,9,50,51).

Some of the genes identified in the present study have previously been associated with more than one disorder and are involved in neurotransmission. This includes *SLC6A4* (identified in the female insula), the molecular target of selective serotonin reuptake inhibitors (SSRIs) which has been associated with sex-dependent effects in depression (52), obsessive-compulsive symptoms (53), and bipolar disorder (54). Another of the genes identified for females in the insula is *GRID2*, a glutamate receptor possibly associated with antipsychotic response (55) which has been associated with obsessive-compulsive symptoms in females but not in males (56). In the hippocampus, a gene only identified in females was *GRIN2A,* a component of the NMDA receptor complex previously associated with schizophrenia (57), bipolar disorder (58) and depression (59). In contrast, one example of neurotransmission-related gene identified as significant only in males is the GABA receptor *GABBR1*, previously associated with schizophrenia, bipolar disorder, depression and autism spectrum disorder (60), and now linked to male thalamic volume. These findings may indicate that sex could modulate neurotransmission pathways, influencing specific brain structure in a sex-dependent manner.

On a different note, females and males may experience various environmental triggers or events throughout their lives (e.g., trauma, pregnancy, parenting), during which their brains undergo adaptations that may have long-lasting effects (61,62). Genetic regulatory factors may modulate these adaptations (63). These include epigenetic modifications and other mechanisms that regulate gene expression – factors that are now recognized as key players in sex variations in human tissues (20,64,65). Our findings appear in line with such evidence since many genes only significant in females or in males were involved in the regulation of gene expression, either through transcriptional regulation (e.g., *FOXP2*, *TP53*, *RFX4*), or through chromatin organization and epigenetic processes (e.g. *BLTP3A*, *ASH1L*, *PHF21A*). In the future, it will be important to elucidate how these regulatory factors interact, respond to environmental triggers, and contribute to the risk of common brain disorders in females and males.

While our results provide important new insights into sex patterns in neurogenetics, they need to be interpreted in the light of some limitations. We studied a population of adults aged 40-80, and the link between genetics, brain and disorder risks may not be the same at all stages of life. Although we minimized the impact of age on our results by carefully matching males and females on age, it will be interesting to explore, in the future, the degree of stability in the observed associations across the lifespan. Lastly, our analyses comparing male and female neurogenetics are descriptive and do not constitute a formal test of sex differences. Instead, they provide a complementary perspective to interaction models by illustrating how genetic signals are distributed across sexes, offering useful context while acknowledging that firm conclusions would require dedicated inferential approaches. By mapping the here identified sex-dependent genes to previous research, our study also highlights avenues for targeted follow-up work with potential relevance for brain disorders.

## Conclusion

In conclusion, while the overall genetic architecture of brain anatomy appears largely similar between females and males, our gene-level results suggest the possibility of sex-dependent features in the genetic underpinnings of brain structure. Future studies are needed to explore sex sensitive neurogenetics across different life stages, and in relation to mental and neurological disorders. Such evidence may improve our understanding of the relationship between sex, genetics and the brain and their role in the onset, presentation, and course of brain disorders, and could also help to identify periods or life events that exacerbate the potential of neurogenetic factors to intensify risk differences between the sexes.

## Declarations

### Acknowledgements

This work was undertaken on the Tjeneste for Sensitive Data (TSD) facilities which is operated/developed by University of Oslo TSD service group (IT-Department), and owned by the University of Oslo as well as on resources provided by the National Infrastructure for High Performance Computing and Data Storage in Norway (UNINETT Sigma2). We would like to thank the UK Biobank team and their research participants, for making this work possible.

### Data availability

Data used in this study were obtained from the UK Biobank (22). GWAS summary statistics and FUMA results will be made openly available upon acceptance.

### Code availability

Codes and software needed to generate the results are available publicly.

MOSTest: https://github.com/precimed/mostest

FUMA: https://fuma.ctglab.nl/

LDSC: https://github.com/bulik/ldsc

