## Supplementary figures and images for "Sex-stratified insights into the genetics of brain volumes in late adulthood"

### Supplementary Figure 1

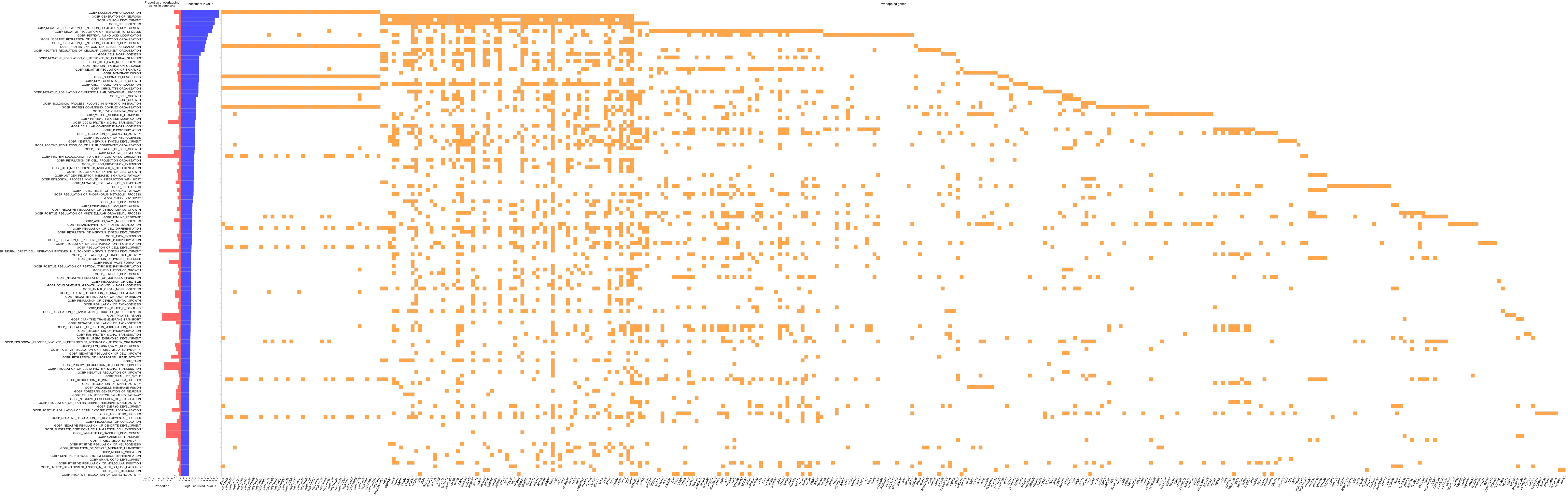

### Supplementary Figure 2

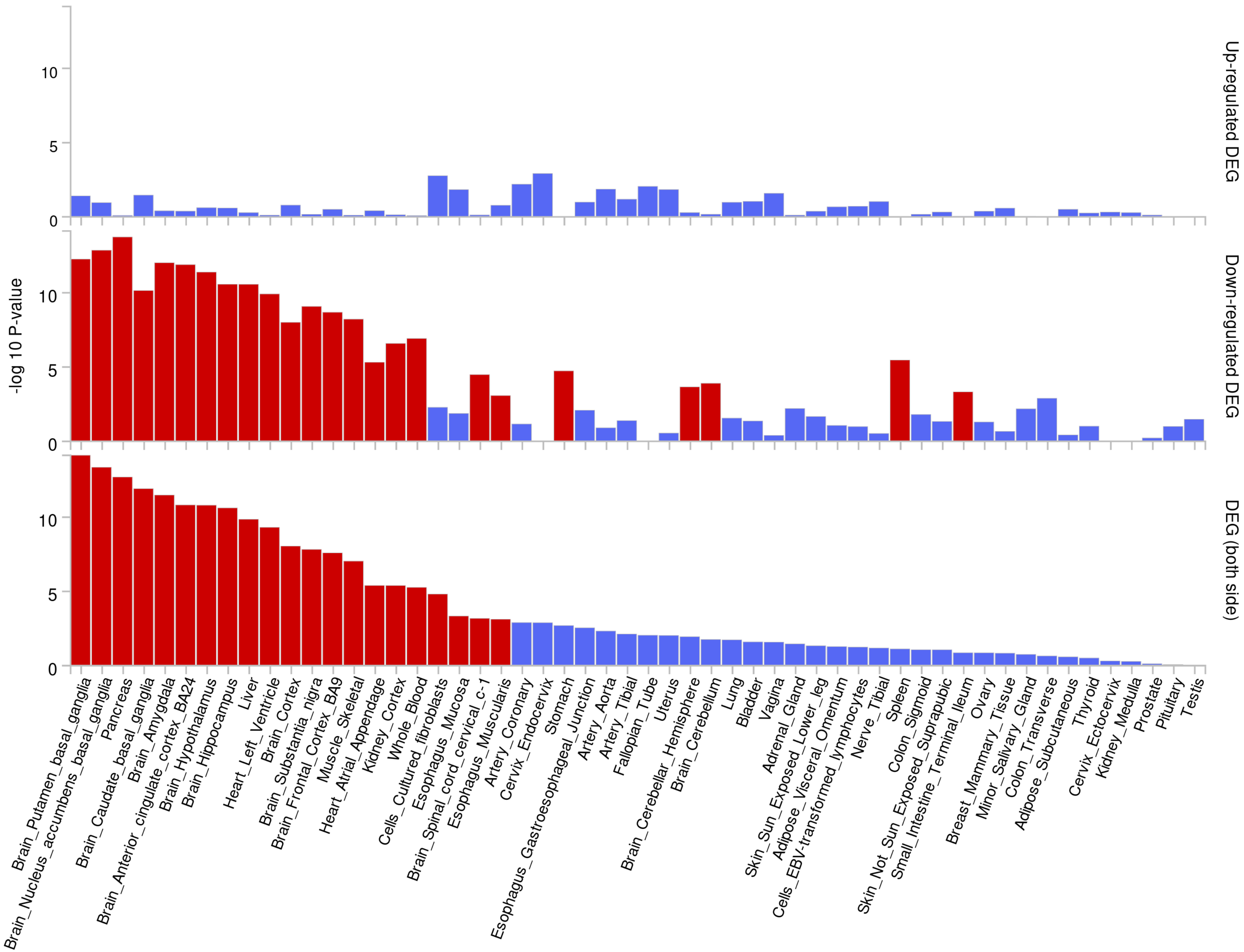

### Supplementary Figure 3

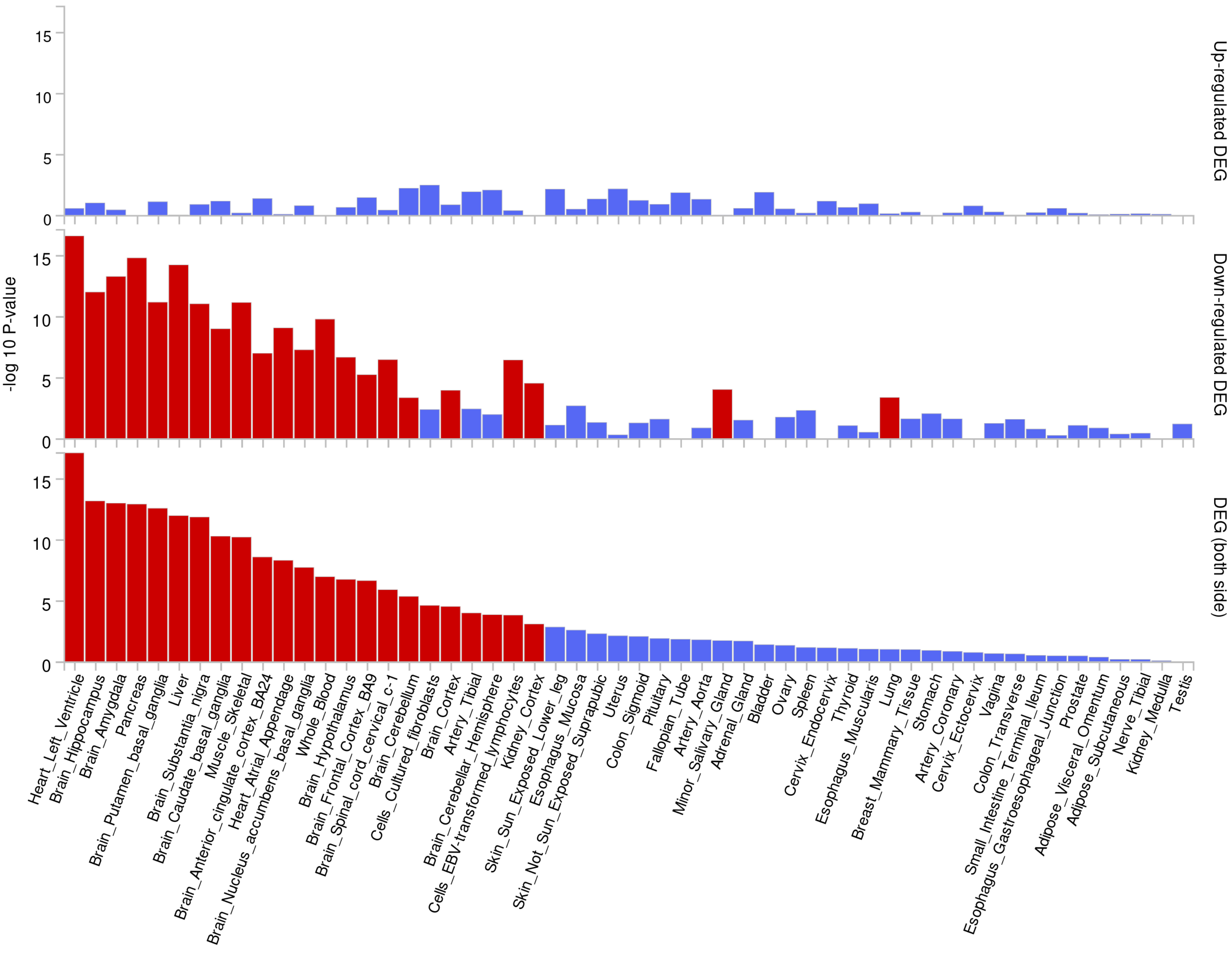

### Supplementary Figure 4

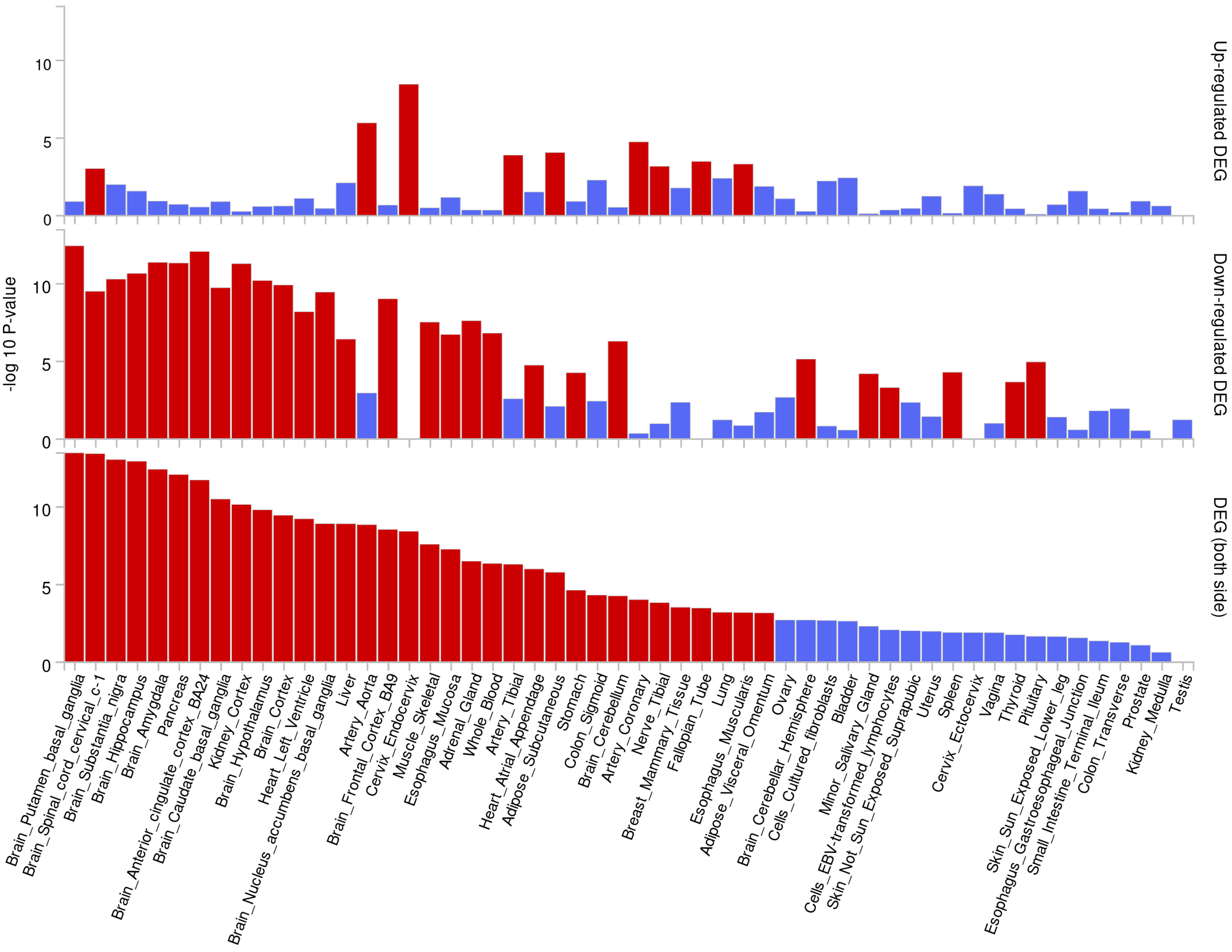

### Supplementary Figure 5

#### A. Females

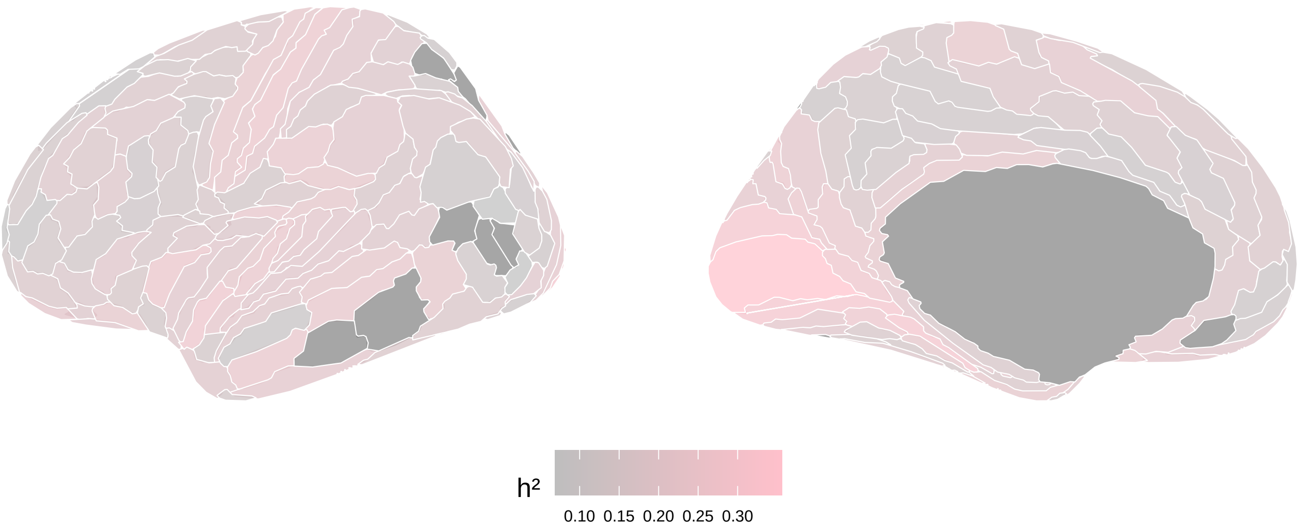


#### B. Males

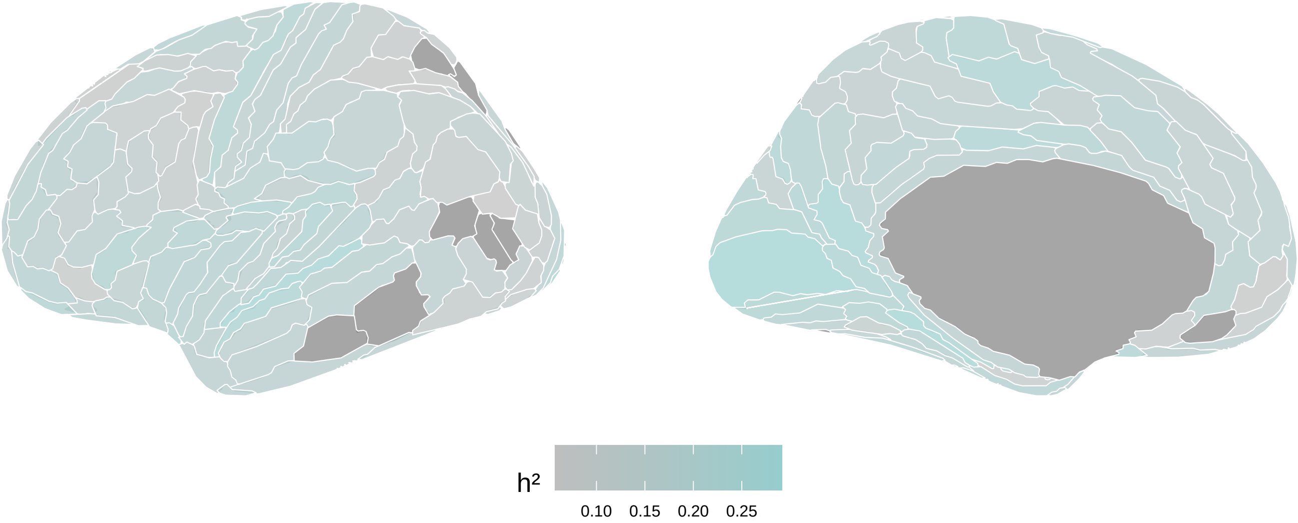
